# Sleep-like changes in neural processing emerge during sleep deprivation in early auditory cortex

**DOI:** 10.1101/2022.03.06.483154

**Authors:** Amit Marmelshtein, Yuval Nir

**Affiliations:** Sagol School of Neuroscience, Tel Aviv University, Tel Aviv 6997801, Israel; Department of Physiology and Pharmacology, Sackler Faculty of Medicine, Tel Aviv University, Tel Aviv 6997801, Israel; Department of Biomedical Engineering, Faculty of Engineering, Tel Aviv University, Tel Aviv 6997801, Israel; The Sieratzki-Sagol Center for Sleep Medicine, Tel Aviv Sourasky Medical Center, Tel Aviv, Israel

**Keywords:** NREM, REM, A1, frequency tuning, rat, click-trains, OFF periods, state-dependent, sensory

## Abstract

Insufficient sleep is commonplace in modern lifestyle and can lead to grave outcomes, yet the changes in neuronal activity accumulating over hours of extended wakefulness remain poorly understood. Specifically, which aspects of cortical processing are affected by sleep deprivation (SD), and whether they also affect early sensory regions, remains unclear. Here, we recorded spiking activity in rat auditory cortex along with polysomnography while presenting sounds during SD followed by recovery sleep. We found that frequency tuning, onset responses, and spontaneous firing rates were largely unaffected by SD. By contrast, SD decreased entrainment to rapid (≥20 Hz) click-trains, increased population synchrony, and increased the prevalence of sleep-like stimulus-induced silent periods, even when ongoing activity was similar. Recovery NREM sleep was associated with similar effects as SD with even greater magnitude, while auditory processing during REM sleep was similar to vigilant wakefulness. Our results show that processes akin to those in NREM sleep invade the activity of cortical circuits during SD, already in early sensory cortex.

## Introduction

Sleep deprivation (SD) is inherent to modern daily life and entails considerable social and health-related costs (Carskadon, 2004). During SD, homeostatic and circadian processes interact to build up sleep pressure (Borbély, 1982) that impairs cognitive performance (Doran et al., 2001), and can lead to serious consequences such as car accidents and medical errors (Carskadon, 2004). Cognitive functions particularly affected by SD include psychomotor and cognitive speed, vigilant and executive attention, working memory, emotional regulation, and higher cognitive abilities (Krause et al., 2017) associated with activity in attentional thalamic and fronto-parietal circuits (Chee et al., 2008; Drummond et al., 1999, 2005; Padilla et al., 2006; Portas et al., 1998; Thomas et al., 2000; Tomasi et al., 2009; Weissman et al., 2006; Wu et al., 2006).

Previous non-invasive studies examined the effect of insufficient sleep on neurophysiological activity (Basner et al., 2013; Chee, 2015; Finelli et al., 2000; Krause et al., 2017; Lorenzo et al., 1995), yet only few studies examined the effects of SD on spiking activities in local neuronal populations. In the rat frontal cortex, robust changes in spontaneous cortical activity gradually emerge during merely a few hours of SD (Vyazovskiy et al., 2011). One study examined the effects of extended wakefulness on sensory responses in high-order human temporal lobe neurons, reporting attenuated, prolonged and delayed responses associated with behavioral lapses (Nir et al., 2017). However, it remains largely unknown whether such effects are restricted to high-order multi-modal regions, or may also affect neuronal activities along specific sensory pathways. Studying the effects of SD on early sensory processing can help shed light on the fundamental processes by which the slow buildup of sleep pressure alters neural processing.

A parallel, equally important, motivation for studying the effects of SD on sensory processing is that it serves as a unique and powerful model for assessing the effects of brain state and arousal on sensory processing at the neuronal level (Harris and Thiele, 2011; Lee and Dan, 2012). A rich body of literature reports the effects of behavioral state and arousal on sensory processing, particularly in the auditory domain. Such studies typically employ one of the following three strategies; One approach is studying how sensory processing differs with respect to behavioral performance on specific tasks (Atiani et al., 2009, 2014; Jaramillo and Zador, 2011; Kato et al., 2015; Otazu et al., 2009). A second approach focuses on momentary changes in arousal, indexed by pupil size, EEG or locomotor activity during wakefulness (Bereshpolova et al., 2011; Lin et al., 2019; McGinley et al., 2015; Zhou et al., 2014; Zhuang et al., 2014). The third strategy contrasts sensory processing in wakefulness with those during anesthesia or natural sleep (Bergman et al., 2022; Issa and Wang, 2011, 2013; Krom et al., 2020; Nir et al., 2013a; Nourski et al., 2018; Raz et al., 2014; Sela et al., 2020). In this context, SD affords an additional unique window to examine how brain states affect sensory processing by offering a ‘middle-tier’ alternative - a state where subjects are awake and responsive but already show behavioral deficits (Krause et al., 2017; Lim and Dinges, 2010). It remains unexplored whether slow accumulation of sleep pressure over hours of SD and extended wakefulness may cause state-dependent changes in sensory processing similar to those associated with momentary arousal changes, on one hand, and to what extent such changes are reminiscent of changes observed during actual sleep, on the other.

Here, we set out to address these issues and examine to what extent SD constitutes an intermediate state between vigilant wakefulness and sleep. We compared neuronal spiking activity in the auditory cortex of freely behaving rats in response to a wide array of sounds including click trains and tones (dynamic random chords (Linden, 2003)). We separately examined how SD affects different aspects of auditory processing including spontaneous activity, frequency tuning, population synchrony, onset vs. sustained responses, and entrainment to slow-vs. fast-varying inputs. Previous research established that some “motifs” of cortical auditory processing are relatively invariant to momentary changes in arousal (e.g. onset responses) whereas other motifs are sensitive to behavioral state (e.g. noise correlations, late sustained responses) (Pachitariu et al., 2015; Sela et al., 2020). Therefore, we hypothesized that cumulative changes over several hours of experimentally-induced SD will lead to changes in *specific* aspects of sensory-evoked activity and that such changes will be detected already in early auditory cortex (Atiani et al., 2009; Jaramillo and Zador, 2011; Otazu et al., 2009; Zhou et al., 2014). In line with this hypothesis, our results show that frequency tuning, onset responses, and spontaneous firing rates were unaffected by SD. By contrast, SD decreased neuronal entrainment to rapid (≥20 Hz) click-trains, increased population synchrony, and increased the prevalence of sleep-like stimulus-induced silent intervals. The changes brought about by SD were qualitatively similar to those observed during recovery NREM sleep, but not during REM sleep where auditory processing was similar to vigilant wakefulness. Thus, our results show that processes akin to those in NREM sleep invade the activity of cortical circuits during SD, already in early sensory cortex.

## Results

To study how sleep deprivation and sleep states affect sensory processing and compare auditory responses across Vigilant, Tired, and sleep conditions, adult male Wister rats (n=7) were implanted with microwire arrays targeting the auditory cortex (AC), as well as EEG and EMG electrodes. After recovery and habituation, rats were placed inside a computer-controlled motorized running wheel within an acoustic chamber for 10 hours starting at light onset (Fig. 1A). We confirmed successful targeting of AC (either A1 or dorsal AC) with histology (Fig. 1B), and by examining the response latency of neuronal units to clicks. 84.9% of recorded units were auditory responsive, and 95.7% of these units (405/423) responded within <20ms (Fig. 1C) attesting to successful targeting of early AC.

**Figure 1.**
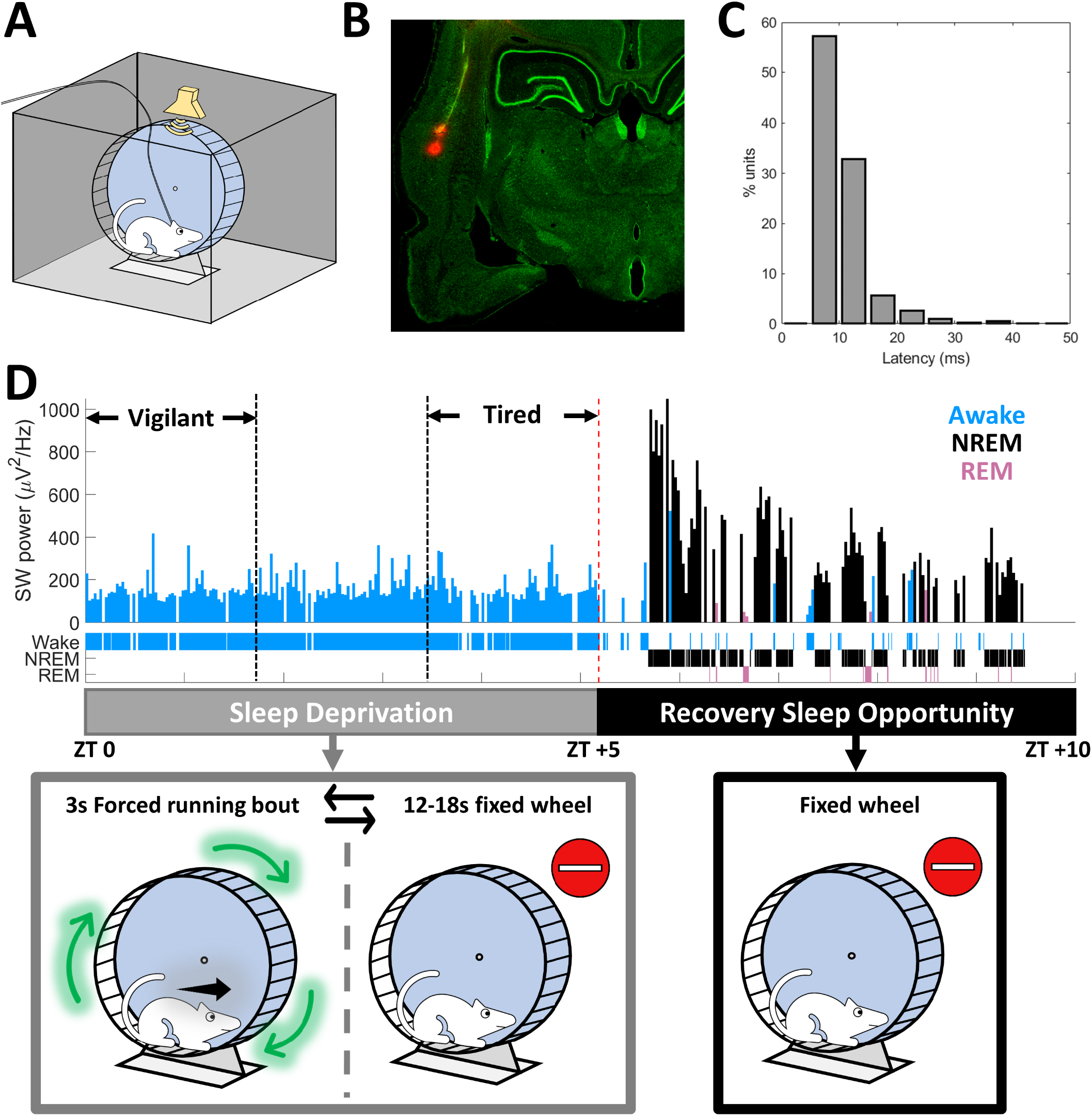
Experimental Setup. A) Experimental setup – Wistar rats were placed inside an acoustic chamber on a motorized running wheel operated intermittently, with an ultrasonic speaker for auditory stimulation and video synchronized with continuous EEG/EMG/intracranial electrophysiology. B) Histology of microwires traces from an array targeting the auditory cortex C) Distribution of response latencies to click stimuli across all responsive units (n=423) attesting to successful micro-electrode targeting to early auditory cortex. D) Top: Representative hypnogram (time-course of sleep/wake states, top) along with dynamics of slow wave activity (SWA, EEG power < 4Hz) in 100s time bins. Bottom: Schematic description of experimental paradigm. Rats were sleep deprived for five hours (zeitgeber time [ZT] 0-5) via intermittent 3s forced running bouts, followed by five hours of recovery sleep opportunity (ZT 5-10), while auditory stimulation was performed continuously throughout the entire experiment with short (∼2s) inter-stimulus-intervals, irrespective of wheel movements.

Rats underwent 5h of sleep deprivation (SD) by intermittent rotations of the wheel (Christie et al., 2008) (3s bouts interleaved with 12-18s idle intervals, Fig. 1D gray). Then, they were left to sleep undisturbed for additional 5h as they spontaneously transitioned between NREM sleep, REM sleep, and short epochs of wakefulness (Fig. 1D, black). Throughout this time, we monitored behavior via synchronized video and intermittently presented auditory stimuli. We focused on comparing the first and last thirds (∼100min each) of the 5h SD period, referred to throughout the manuscript as “Vigilant” and “Tired” conditions, respectively (Fig. 1D). We verified that intervals categorized as Tired were not significantly contaminated by sleep attempts using extensive inspection of video data, and examination of slow wave activity (SWA, 1-4 Hz). Indeed, SWA during the Tired condition was much more similar to that observed in the Vigilant condition than to subsequent NREM sleep (mean±SD: 141±48 *μV*^2^/*Hz* in Vigilant and 184±62 *μV*^2^/*Hz* in Tired, versus 607±218 *μV*^2^/*Hz* in the first third of recovery NREM sleep).

### Frequency tuning, spontaneous firing rates, and onset responses are preserved across Vigilant and Tired conditions during sleep deprivation

Based on previous studies on state-dependent auditory processing (Introduction), we hypothesized that certain features of auditory cortical processing such as frequency tuning will be invariant to SD, whereas other features will be modulated by SD and more generally by arousal state. To test this, we first compared neuronal frequency tuning by examining the responses to dynamic random chord stimuli (Linden, 2003). Fig. 2A shows a representative spectro-temporal receptive field (STRF) of a neuronal cluster during Vigilant and Tired conditions. As can be seen, frequency tuning remains very stable throughout SD. Next, we quantified this stability across the entire dataset (n=198 significantly tuned units out of 496 total) by calculating the tuning width (FWHM, red lines in Fig. 2A) and computing its Modulation Index (MI) across conditions (Fig. 2b, Methods). In line with our hypothesis, we could not reveal a significant change in tuning width across conditions (p=0.529, t_197_=-0.631, Linear Mixed Effects [LME] Model). Indeed, the mean modulation across conditions was -1.85±2.09%, representing only a 1.85% mean decrease in tuning width during the Tired condition. Next, we went beyond tuning width and examined more generally whether the frequency tuning profile of each neuron is stable across states, representing additional features such as preferred frequency and temporal dynamics of tuned responses. To this end, we calculated the signal correlation between STRF maps in Vigilant and Tired conditions (Fig. 2C middle). We then compared it with signal correlation benchmarks for minimal correlation (Fig. 2C left: different units in different conditions) and maximum correlation (Fig. 2C right: same units, between 1^st^ and 2^nd^ halves of data within the same condition, Methods). We found that the STRF signal correlation between Vigilant and Tired conditions (middle bar, 0.638±0.012) was significantly higher than between different units (left bar, 0.219±0.009, p=4.53 × 10^−18^, t_197_=9.57, LME), and virtually as high as the signal correlation within each condition (middle vs. right bar, 0.638±0.012 vs. 0.647±0.012; p=0.16, t_197_=-1.41, LME). Given a finite number of trials and some inevitable degree noise in the data, STRF profiles across Vigilant and Tired conditions are as similar as they possibly can be. Thus, both tuning width and the signal correlation of STRF profiles were invariant to changes in arousal states during sleep deprivation.

**Figure 2.**
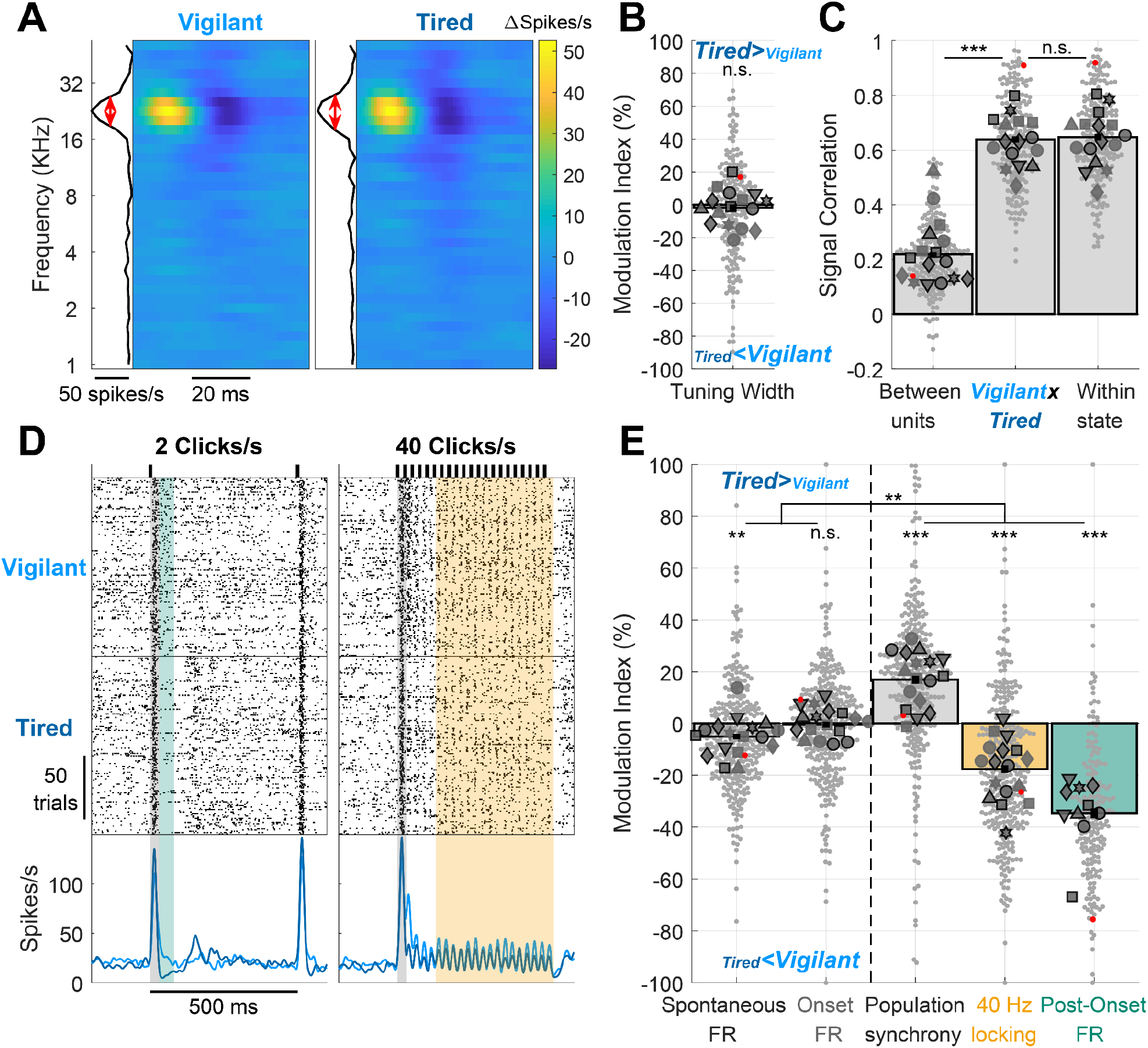
Auditory cortex processing during sleep deprivation. A) Representative spectro-temporal receptive field (STRF) of a unit in auditory cortex showing preserved frequency tuning across Vigilant and Tired conditions (left and right, respectively). B) Modulation of frequency tuning width (Tired vs. Vigilant conditions) for all tuned units (n=198 out of 496 total) and sessions (n=16). C) Signal correlations of frequency tuning across the entire dataset between different units in the same session (left bar, benchmark for min. correlation), between Vigilant and Tired conditions of the same individual units (middle bar) and between 1st and 2nd halves of trials in the same condition for the same individual units (right bar, benchmark for max. correlation). Note that signal correlations are nearly as high across Vigilant and Tired conditions as they are within the same condition. D) Representative raster and peri-stimulus time histogram (PSTH) for a unit in response to 2 and 40 clicks/s click trains (left and right, respectively). Gray shading marks the onset response [0-30]ms period. Green shading represents the post-onset [30-80]ms period where firing rate was especially attenuated during the Tired condition. Yellow shading represents the [130-530]ms period where sustained locking to the 40 click/s train was attenuated during the Tired condition. E) Modulation of activity/response features between Tired and Vigilant conditions across units (n=327) and sessions (n=17). Features (left to right) denote: spontaneous firing rate (FR), onset response FR, population synchrony, 40-Hz locking and post onset FR. 2 click/s train were presented in 11 out of 19 sessions (‘auditory paradigm A’, n=199 units). For Panels B, C and E small gray markers represent individual units. large dark gray markers represent mean of all units in an individual session. Each marker shape represents sessions from an individual animal. Markers with/without black edges represent ‘auditory paradigm A’ and ‘auditory paradigm B’ sessions, respectively. Red dots point to the representative unit presented in panels A and D. Dashed vertical line separates features minimally/not significantly affected by condition (spontaneous FR and onset response FR; on left) vs. features significantly that are disrupted in the Tired condition (population synchrony, 40Hz locking, and post-onset FR; on right).

We proceeded to analyze neuronal responses to 500ms click trains at different rates (Fig. 2D, 2, 10, 20 & 30 clicks/s in 11 experimental sessions and 40 clicks/s in 19 experimental sessions). We first quantified onset response magnitude to the 40 clicks/s stimulus, as well as spontaneous (baseline) firing rate preceding stimulus onset across all responsive units (65.3%, 324 of 496 units) in Vigilant and Tired conditions. As can be seen in a representative unit (Fig. 2D), the spontaneous firing rate did not change between conditions. Similarly, the onset response (gray shading, [0-30]ms relative to stimulus onset) was similar in magnitude across conditions. Quantitative analysis across the entire dataset (Fig 2E, two leftmost bars) revealed a slight reduction (−5.1±1.1%) in spontaneous firing during the Tired condition (p=0.0085, t_323_=-2.65, LME), while onset FR did not exhibit significant modulation (−0.52±1.17%, p= 0.92, t_323_=0.1, LME). Overall, some aspects of cortical auditory processing, including frequency tuning, spontaneous firing, and onset responses are largely preserved across Vigilant and Tired conditions during sleep deprivation.

### Population synchrony, entrainment to fast click-trains, and post-onset silence are strongly modulated by sleep deprivation

Next, we tested the degree to which sleep deprivation affects other features of cortical auditory processing. We predicted that population synchrony would increase in Tired condition given the increased propensity of local neuronal populations to exhibit synchronous OFF-states in SD (Vyazovskiy et al., 2011). Quantifying “population coupling” (Okun et al., 2015), a measure of how correlated each unit’s firing is with the firing of the local population, we found a significant increase (17±1.6%) in population synchrony during the Tired condition (Fig. 2E, p=5.2 × 10^−10^, t_323_=6.41, LME). We also predicted that entrainment to fast click trains (40 clicks/s) might be especially sensitive to sleep deprivation (Krom et al., 2020; Plourde, 1996; Sharon and Nir, 2018). As can be seen in a representative example (Fig. 2D, orange shading), the magnitude of sustained locking to the click train decreased during the Tired condition. A quantitative analysis across the entire dataset revealed a significant decrease of 17.7±1.5% in 40Hz Locking (Fig. 2E orange bar, p=1.4×10^−6^, t_323_=-4.92, LME).

When presenting click trains at slower rates (2 & 10 clicks/s, n=11 sessions), we observed that the onset response ([0,30]ms) was followed by a post-onset period ([30,80]ms) exhibiting robust firing attenuation in the Tired condition (Fig. 2D left, green shading). Indeed, post-onset firing was significantly attenuated in the Tired condition compared to the Vigilant condition (34.6±1.9%, Fig. 2E green bar, p=4.6×10^−14^, t_195_=-8.14, LME). Post-onset firing reduction emerged as a particularly state-sensitive aspect of the cortical auditory response, showing significantly stronger modulation than population synchrony and 40Hz locking (p≤0.0151, df=195, LME). Analysis of variance among the distinct features of cortical auditory processing confirmed that they are differentially modulated by SD (p=2.8×10^−4^, n=7 animals, Friedman test). Pair-wise comparisons revealed that while spontaneous firing rates and onset responses were largely preserved, population synchrony, 40-Hz click train locking, and post-onset firing were modulated significantly more strongly than the former two features (p≤0.0025, df=323 or 195, LME). In addition, an hour-by-hour analysis revealed that auditory processing features that were sensitive to SD exhibited gradually accumulating changes, corresponding to gradually accumulating sleep pressure (Supp. Fig. 1).

### The effects of sleep deprivation on cortical auditory processing mimic those of NREM sleep

Previous studies have shown that during Tired conditions upon SD, features of NREM sleep activity (e.g. slow/theta activities and OFF-states) ‘invade’ the ongoing activity of cortical circuits (Finelli et al., 2000; Nir et al., 2017; Vyazovskiy et al., 2011). We wondered if the same is true for stimulus-driven activity, and whether it can already be observed in early sensory cortex. To test this, we compared neural activity and auditory responses during the Vigilant condition with those during the 5h recovery sleep period (Fig. 3), when rats spent 48±7.5% of time in NREM sleep, 22±7.6% of time in wakefulness, and 6.5±4.2% of time in REM sleep (mean±SD, additional intervals in transition or undetermined states, not analyzed further). We hypothesized that features showing similarity across Vigilant and Tired conditions will also be invariant to full-fledged NREM sleep, whereas changes observed during SD will be accentuated in recovery sleep data.

**Figure 3.**
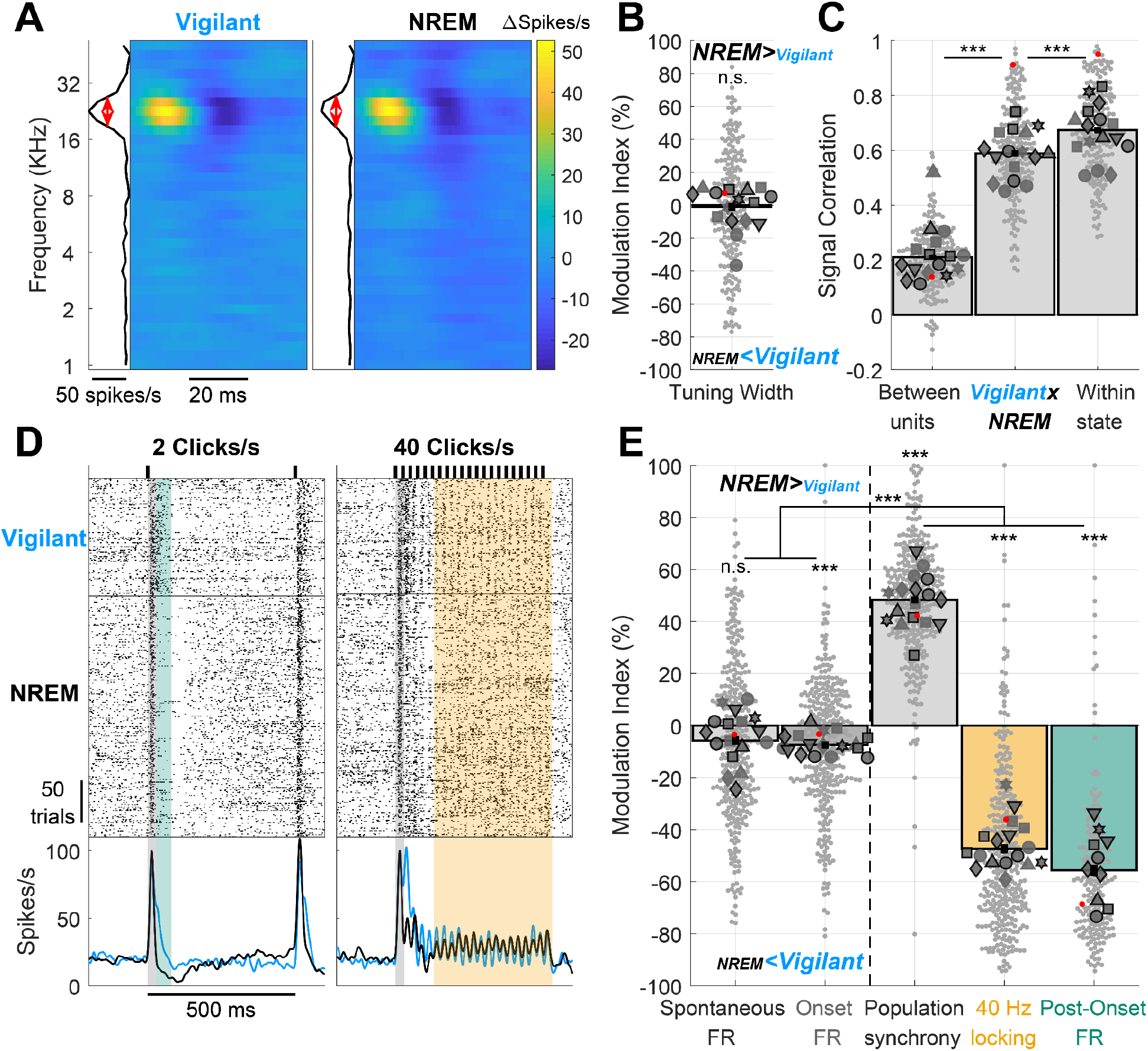
Auditory cortex processing during recovery NREM sleep vs. vigilant wakefulness. Same as Fig. 2 but comparing recovery NREM sleep to the Vigilant condition. A) Representative spectro-temporal receptive field (STRF) of a unit in auditory cortex showing preserved frequency tuning across Vigilant and NREM sleep conditions (left and right, respectively). B) Modulation of frequency tuning width (NREM sleep vs. Vigilant conditions) for all units (n=200) and sessions (n=16). C) Signal correlations of frequency tuning across the entire dataset between different units in the same session (left bar), between Vigilant and NREM sleep conditions of the same individual units (middle bar) and between 1st and 2nd halves of trials in the same condition for the same individual units (right bar). Note that signal correlations are nearly as high across Vigilant and NREM sleep conditions as they are within the same condition. D) Representative raster and peri-stimulus time histogram (PSTH) for a unit in response to 2 and 40 clicks/s click trains (left and right, respectively). Gray shading marks the onset response [0-30]ms period. Green shading represents the post-onset [30-80]ms period where firing rate was especially attenuated during the Tired condition. Yellow shading represents the [130-530]ms period where sustained locking to the 40 click/s train was attenuated during the NREM sleep condition. E) Modulation of activity/response features between NREM sleep and Vigilant conditions across units (n=327) and sessions (n=17). Features (left to right) denote: spontaneous firing rate (FR), onset response FR, population synchrony, 40-Hz locking and post onset FR. 2 click/s train were presented in 11 out of 19 sessions (‘auditory paradigm A’, green bar, n=199 units, 10 session). For Panels B,C,E, small gray markers represent individual units. large dark gray markers represent mean of all units in an individual session. Each marker shape represents sessions from an individual animal. Markers with/without black edges represent ‘auditory paradigm A’ and ‘auditory paradigm B’ sessions, respectively. Red dots point to the representative unit presented in panels A and D. Dashed vertical line separates features minimally/not significantly affected by condition (spontaneous FR and onset response FR; on left) vs. features that are significantly disrupted in the NREM sleep condition (population synchrony, 40Hz locking, and post-onset FR; on right).

Indeed, frequency tuning, spontaneous firing, and onset responses were similar across Vigilant and NREM sleep conditions. Fig. 3A shows an example STRF during Vigilant and NREM sleep conditions. As observed during SD, full-fledged NREM sleep did not alter neuronal frequency tuning. Across the entire dataset, mean frequency tuning width did not significantly change (−1.18±1.25%, p=0.642, t_197_=-0.47, LME), and the STRF profile signal correlation (Fig. 3D) between Vigilant and NREM sleep conditions (middle bar) was nearly as high as the signal correlation within each condition (right bar). Although the difference in signal correlation was highly significant statistically (p=3.5×10^−11^, t_197_=-7.02, LME), its magnitude was moderate: signal correlation between Vigilant and NREM sleep was 87.4% of the mean signal correlation within each condition (0.589 vs. 0.674).

Fig. 3D shows that for this representative unit, spontaneous firing and onset responses were also unchanged during NREM sleep, contrasting with strong modulation of post-onset firing and 40Hz-locking (green and yellow shading, respectively). Analysis of the entire dataset confirmed a modest attenuation of spontaneous firing and onset responses during recovery NREM sleep (Fig 3E left, 5.78±1.68% and 7.45±1.38%, respectively), that was statistically significant only for onset responses (p=1.1×10^−6^, t_323_=-4.98 for onset response and p=0.16, t_323_=-1.39 for spontaneous FR, LME). In sharp contrast, these modest changes were overshadowed by strong modulations of population synchrony (48.3±1.3%), 40 Hz Locking (47.4±1.73%) and post-onset firing (55.7±2.19%) during NREM sleep (Fig. 3E right, p≤1.2 × 10^−27^). As was the case for SD, the differential modulation of specific features of cortical auditory processing by NREM sleep was highly significant (p=1.7×10^−4^, n=7 animals, Friedman test) where population synchrony, 40Hz-locking, and post-onset firing were significantly more modulated than spontaneous firing and onset responses (p≤2.8×10^−16^ for all pair-wise comparisons, df=323 or 195, LME). Overall, the same aspects of cortical auditory processing that showed maximal modulation during SD (population synchrony, 40Hz Locking, post onset FR) were maximally modulated during recovery NREM sleep.

### Sleep deprivation and NREM sleep entail sensory adaptation at lower frequencies

To better understand how Tired and NREM sleep states disrupt locking to click trains, we presented click rates at various rates (2, 10, 20, 30 and 40 clicks/s, n=11 sessions). As can be seen in a representative response (Fig. 4A), sustained locking to slower click trains (2 and 10 clicks/s, yellow shading) was stable during the Tired and NREM sleep conditions relative to Vigilant. In contrast, locking to faster click trains (≥20 clicks/s) showed strong attenuation. We thus quantified the modulation in response locking across the entire dataset during SD (Tired vs. Vigilant, 194 units, Fig 4B) and during NREM sleep (NREM vs. Vigilant, Fig 4C). Locking to different click rates was differentially modulated by SD (Fig. 4B, p=9.7×10^−4^, n=6 animals, Friedman test). Pairwise comparisons revealed that locking to faster click-trains (20, 30 & 40 clicks/s) was significantly more attenuated than to slower click trains (2 & 10 Clicks/s), with an average attenuation of 18.4% vs. 4.4%, respectively (for all comparisons p≤2.6×10^−4^, df=193, LME, mean MI: - 3.05±1.46%, -5.71±1.36%, -17.2±1.75%, -19.1±1.67% and -18.9±1.87%, for 2, 10, 20, 30 and 40 clicks/s, respectively). NREM sleep showed qualitatively similar and stronger effects (Fig. 4C, p=0.0036, n=6 animals, Friedman test, mean MI: -11.6±1.93%, -31.2±1.99%, -43.7±1.84%, -50.9±1.98% and -49±2.05%, for 2, 10, 20, 30 and 40 clicks/s, respectively). Pairwise comparisons revealed a gradual modulation during NREM sleep depending on click-train rate (−11.6% for 2 click/s versus -31.2% for 10 clicks/s, and even stronger attenuations for 20, 30 & 40 clicks/s, p≤0.0057, df=193, for all comparisons, LME).

**Figure 4.**
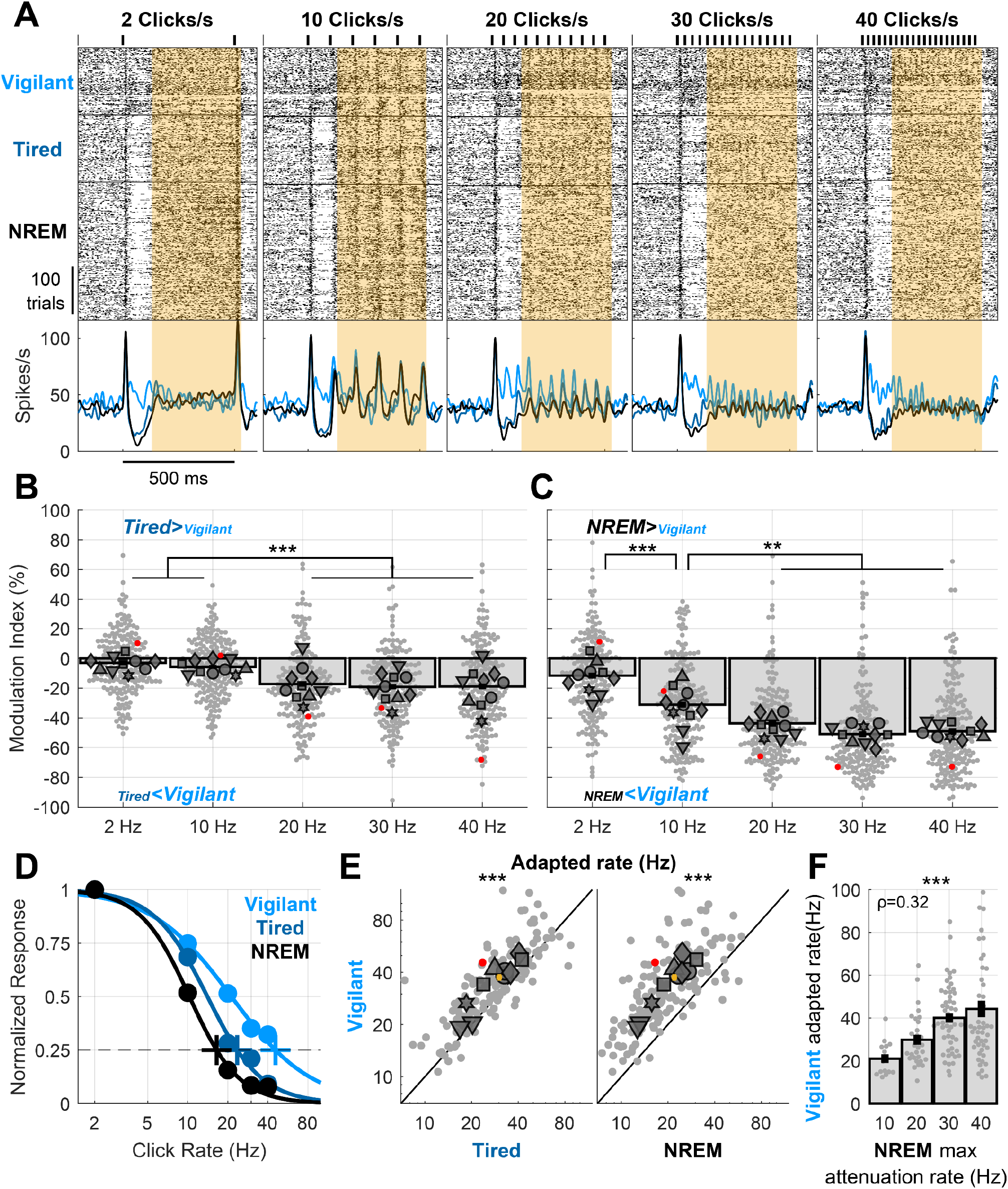
Recovery NREM sleep and sleep deprivation both entail a shift in sensory adaptation. A) Representative unit raster and PSTH responses to 2, 10, 20, 30 and 40 clicks/s responses. Note that locking to click trains is progressively more disrupted in Tired/NREM sleep conditions with increasing click train rate. B) Modulation of locking to different click rates (Tired vs. Vigilant) for all units (n=197) and sessions (n=10). Locking to fast click trains (≥20 clicks/s) is significantly attenuated during sleep deprivation (‘Tired’). C) Same as B but comparing recovery NREM sleep to the Vigilant condition, showing increasingly stronger attenuation for faster click trains. D) Normalized locked responses in a representative unit (y-axis) as a function of click rate (x-axis) separately for Vigilant (cyan), Tired (blue), and recovery NREM sleep (green) conditions. Circles represent the observed locked response to each click rate in each condition. Thick traces connecting the circles represent the best sigmoid fit. Cross represents the estimated ‘adapted click-rate’, i.e. the click rate for which the normalized response would be 25% of maximum. E) Left: scatter plot of the ‘adapted click-rate’ for all units and sessions, comparing Vigilant (y-axis) with Tired conditions (y-axis); Right: same when comparing Vigilant (y-axis) with recovery NREM sleep (x-axis). Yellow cross represents mean±SEM across all units (n=150). F) observed click rate for which units demonstrate maximal attenuation between Vigilant and NREM sleep conditions (x-axis, Methods) vs. the estimated ‘adapted click-rate’ during the Vigilant condition (y-axis). Note that units with lower ‘adapted click-rate’ during wakefulness also show lower attenuation rates when comparing NREM sleep vs. Vigilant. For Panels B,C,E,F: small gray markers represent individual units. Large dark gray markers represent mean of all units in an individual session. Red dots point to the representative unit presented in panels A and D.

To capture how different arousal states affect the entire sensory adaptation curve, we fitted a sigmoid function to describe how response attenuation changes with increasing click rate (Methods). Fig 4D shows this fit for the same unit example shown in Fig. 4A. We then estimated the *“adapted rate”*, i.e. the click rate for which the response is attenuated to 25% of its maximum (Methods), for each arousal condition separately (crosses in Fig 4D). For the example unit shown, the estimated *adapted rate* during the Vigilant condition was 45.6 clicks/s, decreased to 23 clicks/s during the Tired condition, and decreased further to 16.6 clicks/s during NREM sleep. A quantitative analysis across the entire dataset (Fig. 4E) revealed that SD decreased the adapted rate by 15.6±1.84% (Fig. 4E left, p=1.1×10^−7^, t_146_=-5.58, LME, Vigilant: 32.9 vs. Tired: 26.8 clicks/s). An even stronger decrease of 36.3±1.87% was observed in NREM sleep (Fig. 4E right, p=3.9×10^−36^, t_146_=-16.9, LME, Vigilant: 32.9 vs. NREM sleep: 19.7 clicks/s). Overall, Tired and NREM sleep low-arousal states shift the sensory adaptation gain curve to lower frequencies.

Could it be that some neurons are strongly adapted to begin with, and these are the neurons who are most sensitive to changes in state? To examine this, we tested whether neurons that show a low adapted rate during the Vigilant condition (e.g. weak locking already for 10 click/s) may correspondingly show a strong attenuation at lower frequencies during NREM sleep (compared to the Vigilant condition). We calculated for each neuron its estimated ‘*adapted rate*’ during the Vigilant condition, and compared it to the click rate showing maximal attenuation during NREM sleep (Fig. 4F, Methods). For example, the representative unit in Fig. 4A,D shows a close-to-maximal attenuation during NREM sleep already at 20 clicks/s (red points in Fig. 4C), while its estimated ‘adapted rate’ during the Vigilant condition was 45.6 clicks/s (light blue cross in Fig. 4D). Analysis across the entire dataset confirmed the significant correlation between these two measures (Fig. 4F, p=6.2×10^−5^, rho=0.324, for n=147 units, Spearman Correlation). Such correlation was not significant when comparing Vigilant and Tired conditions (p=0.46, rho=0.062), possibly due to the weaker modulation observed in SD. Thus, we found that for a given neuron, the attenuation during NREM sleep is dictated by the sensory adaptation curve during vigilance, such that neurons showing significantly adapted response at lower click rates are also attenuated during NREM at lower click rates. Finally, linear modeling revealed that reduced locking to fast click trains cannot be simply explained by post-onset reduction in firing (Supp. Fig 2).

### Stimulus-induced silent intervals are more sensitive to sleep deprivation than spontaneous silences

The most sensitive measure of cortical auditory processing in low-arousal states was a reduction or complete cessation of firing following the onset response in Tired and NREM sleep conditions (Fig. 5A Left, [30,80]ms post click, green shading). Given that such stimulus-induced silence was reminiscent of an ‘OFF-state’ observed in ongoing activity of local neuronal populations during NREM sleep and SD (Vyazovskiy et al., 2011), we examined if it likewise represents a network-wide event or, alternatively, simply reflects a refractory-like period in spiking of individual neurons exposed by the onset response to the auditory stimulus. To test this, we compared each auditory trial (with its stimulus-induced onset response and post-onset silence, Fig 5A, left) with a matched interval of ongoing activity containing similar spiking bursts (Fig. 5A, right). We found that post-onset FR reduction was only apparent following auditory stimulation and onset responses but not present in spontaneous firing (Fig. 5A,B, green shading): baseline-normalized post onset FR was gradually reduced from 0.98±0.039 during the Vigilant condition to 0.63±0.033 during the Tired condition, and even further to 0.38±0.023 during NREM sleep (Fig. 5C, p<3 × 10^−7^ for all pair-wise comparisons, df=195, LME). By contrast, FR following spontaneous bursts only revealed marginal changes across conditions: Vigilant: 1.08±0.016, Tired:1.01±0.016, NREM: 0.98±0.02 (p=0.032 for comparing Vigilant and Tired conditions, p>0.05 for all other comparisons, df=195, LME). The attenuation in click-induced post-onset FR during the Tired condition (34.2±1.95% relative to vigilance) was significantly larger than that following spontaneous-bursts (5.2±1.7%, p=1.5 × 10^−18^, t_195_=9.75, LME), as was also true for NREM sleep (p=1.5 × 10^−16^, t_195_=9.04, LME). Thus, post-onset suppression isn’t simply a property of individual neurons that reduce firing after vigorous activity, but represents a network event induced by the stimulus.

**Figure 5.**
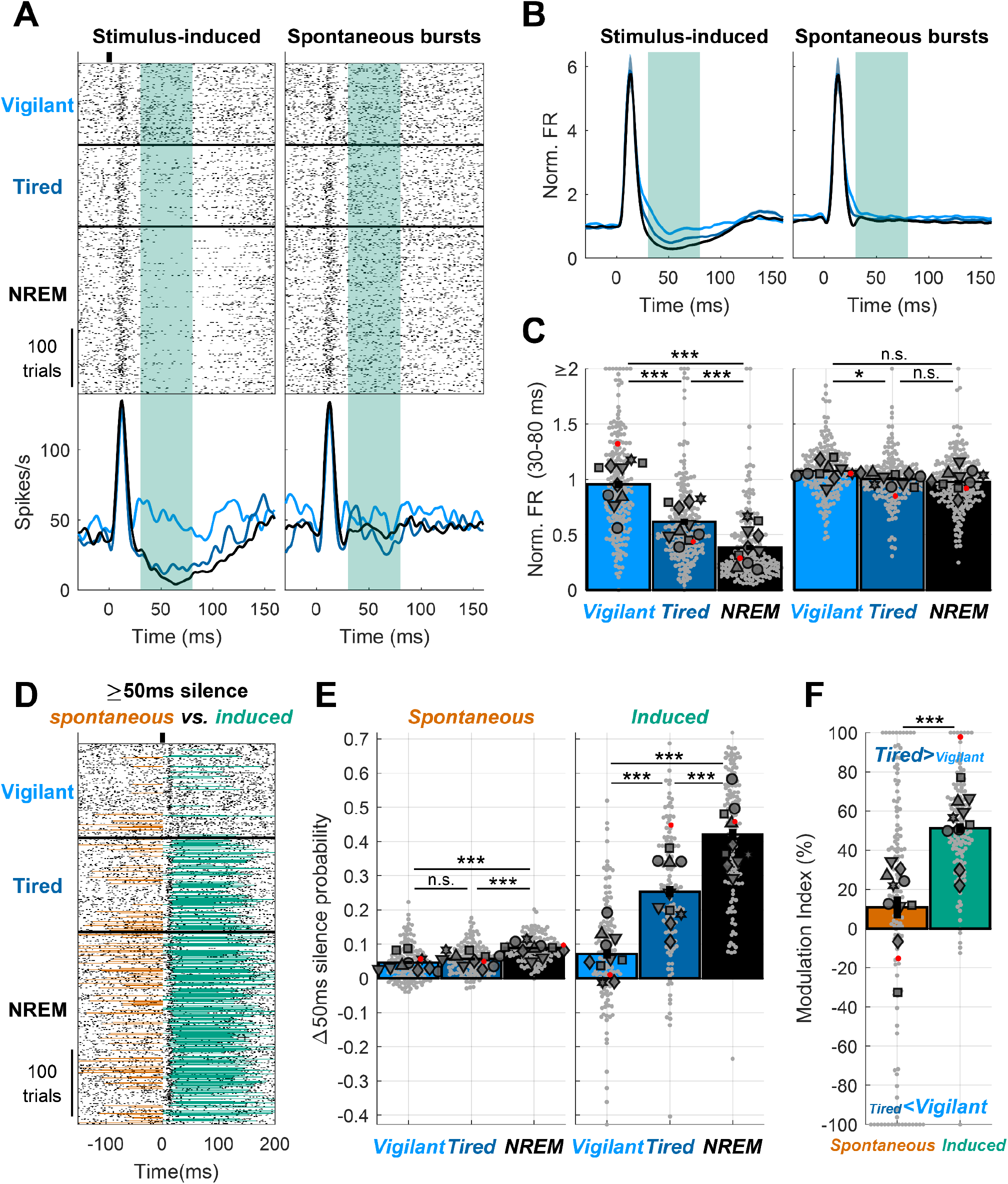
Stimulus-induced silent intervals are especially sensitive to sleep deprivation. A) Representative unit raster and PSTH response to a click across Vigilant, Tired and NREM sleep conditions (left) and trial-by-trial matched, equally strong, spontaneous bursts (matching the [0,30]ms click onset response) of the same unit (right). Note that there is no post-onset FR reduction following spontaneous bursts. Green shading represents the post-onset [30,80]ms period. B) mean normalized PSTH of all units (n=195) for the stimulus(click)-induced response (left), and matched spontaneous bursts (right) across Vigilant, Tired and NREM sleep conditions. C) Post-onset normalized FR across Vigilant, Tired and NREM sleep conditions for the stimulus-induced response (left) and the matched spontaneous bursts (right) for all units (n=195) and sessions (n=10). D) Representative unit raster and PSTH response to 2 clicks/s train across Vigilant, Tired and NREM sleep conditions. Silent intervals (≥50ms firing silence) just preceding ([-50,0]ms) or immediately following ([30,80]ms) stimulus onset are marked in orange and green, respectively. Note that spontaneous silent intervals (orange) are prevalent in NREM sleep but rare during the Tired condition (as in Vigilant), whereas stimulus-induced silent intervals (green) strongly increase in the Tired condition. E) Increase in silent intervals probability (relative to Poisson process with the same spontaneous firing rate) across Vigilant, Tired and NREM sleep conditions, separately for spontaneous (left) and stimulus-induced (right) silent intervals for all electrodes (n=126) and sessions (n=10). F) Modulation of the probability of spontaneous and stimulus-induced silent intervals across Tired vs. Vigilant conditions for all electrodes (n=126) and sessions (n=10). Stimulus-induced silent intervals show a larger and more reliable change upon sleep deprivation (comparing Tired and Vigilant conditions) relative to spontaneous intervals. Bars represent mean across all units/channels. Small gray markers represent individual units/channels. Large dark gray markers represent mean of all units/channels in an individual session. Red dots point to the representative unit presented in panels A and D.

Next, we complemented the analysis of graded firing rate reductions with a binary approach of detecting OFF periods – intervals of neuronal silence ≥50ms, typically observed in ongoing sleep activity and in SD. Both spontaneous and stimulus-induced silent intervals (presumably OFF-states) were rare during the Vigilant condition but more frequent during NREM sleep (Fig. 5D). During the Tired condition (wakefulness after several hours of SD), stimulus-induced silent intervals were very frequent while spontaneous silent intervals continued to be rare. We analyzed the probability of spontaneous silent intervals relative to a random Poisson process (Fig. 5E, Methods) across the entire dataset, and found a graded modulation by arousal state (Vigilant: 4.61±0.48%, Tired: 5.67±0.38%, NREM sleep: 8.87±0.33%, p=0.0057, n=6 animals, Friedman test). Pair-wise comparisons revealed that the probability in NREM sleep was significantly greater than other conditions (p≤9.7 × 10^−6^, df=126, LME, compared to Vigilant and Tired conditions), while the increase from Vigilant to the Tired condition exhibited a non-significant trend (p=0.0501, t_125_=-1.98, LME). By contrast to spontaneous silent intervals, the probability of stimulus-induced silent intervals was higher and more strongly modulated by condition (Vigilant: 7.14±1.38%p, Tired: 25.4±1.63%, NREM sleep = 42±1.6%, p=0.0025, n=6 animals, Friedman test, Fig. 5B right, p≤4.1×10^−11^, df=125, for all pairwise comparisons, LME). In the Vigilant condition, the probability of stimulus-induced silent intervals was not significantly different than that of spontaneous silent intervals (p=0.236, t_125_=1.19, LME, Spontaneous: 4.61±0.48% vs. Induced: 7.14±1.38%) but this difference was highly significant in the Tired and NREM sleep conditions (p≤9×10^−8^, df=125, LME, for all comparisons, Spontaneous: 5.67±0.38%, 8.87±0.33%, vs. Induced: 25.4±1.63%, 42±1.6% for Tired and NREM sleep, respectively). Indeed, the mean modulation index comparing silent interval probability in Tired vs. Vigilant conditions (Fig. 5F) increased significantly from 10.8±5.55% for spontaneous silent intervals to 51.2±2.5% for stimulus-induced silent intervals (p=5.4 × 10^−5^, t_125_=4.18, LME). Overall, these results establish that stimulus-induced silent intervals reveal a hidden facet of neural processing during SD that goes beyond what is observed in spontaneous activity.

### Auditory processing during REM sleep resembles the Vigilant condition, unlike NREM sleep

REM sleep is a unique (‘paradoxical’) behavioral state that is characterized both by disengagement from the environment co-occurring with desynchronized cortical activity and often accompanied by vivid dreams(Nir and Tononi, 2010). Therefore, REM offers a unique lens through which to examine the changes in cortical auditory processing, potentially revealing which aspects are similar to NREM sleep (likely reflecting a general feature of sleep and sensory disengagement) and which aspects are similar to the Vigilant condition (likely reflecting a general feature of desynchronized cortical states).

We first observed that frequency tuning was stable during REM sleep (Fig. 6A). Across the entire dataset, tuning width was reduced by an average of 15% (Fig. 6B, p=3.3×10^−4^, t_120_=-3.7, LME), while the signal correlation between the Vigilant and REM sleep conditions (Fig. 6C middle bar, 0.547±0.014) was nearly as high (89.7%) as the maximal benchmark within each condition (Fig 6C, 0.609±0.014, p=5.5×10^−8^, t_120_=-5.8, LME). Next, examining different aspects of the neuronal activity and auditory response revealed that REM sleep exhibits a very similar profile to the Vigilant condition (Fig. 6D,E). Unlike NREM sleep, REM sleep was associated with high post-onset firing and strong locking to the 40 click/s train, as in the Vigilant condition. A quantitative analysis across the entire dataset revealed modest average difference between REM sleep and the Vigilant condition (all mean MI<21%, Fig. 6E). Moreover, in measures such as spontaneous firing and 40Hz-locking, REM sleep was even significantly higher than the Vigilant condition (Fig. 6E, all p<4.3×10^−8^, df=322, LME). Conversely, when contrasting REM sleep with NREM sleep, strong and reliable differences emerged (Fig. 6F). As was the case when comparing Vigilant condition with NREM sleep, different aspects of the neuronal activity and auditory response were differentially modulated by state (Fig. 6F, p=4.1 × 10^−4^, df=7 animals, Friedman test). Again, the onset response was minimally affected by the state (MI: 10.3±1.32%, p=1.9×10^−13^, t_323_=7.68, LME). Spontaneous firing increased by 26.8±1.4% during REM sleep compared to NREM sleep (p=2.7×10^−21^, t_323_=10.2, LME). Even larger changes were observed when comparing population synchrony (MI: -37.5±1.52%, p=1.04×10^−16^, t_323_=-8.77, LME), 40-Hz locking (MI: 61.7±1.35%, p=2.6×10^−91^, t_323_=28.8, LME) and post onset firing (MI: 64.4±1.95%, p=1.5×10^−11^, t_195_=7.17, LME). The *‘adapted rate’* (Fig. 6G) during REM sleep was higher than during the Vigilant condition, and increased on average from 31.8 to 38.8 clicks/s (Fig. 6H left, p=3.4×10^−7^, t_135_=5.37, LME). Conversely, robust differences in the adapted rate emerged when comparing REM sleep (38.8 clicks/s) to NREM sleep (19.5 clicks/s; Fig. 6H right, p=7.1×10^−46^, t_135_=21.7, LME). Altogether, cortical auditory processing during REM sleep is dramatically different from that in NREM sleep, showing a profile similar to that observed during the Vigilant condition (and in some aspects exhibiting even stronger activity).

**Figure 6.**
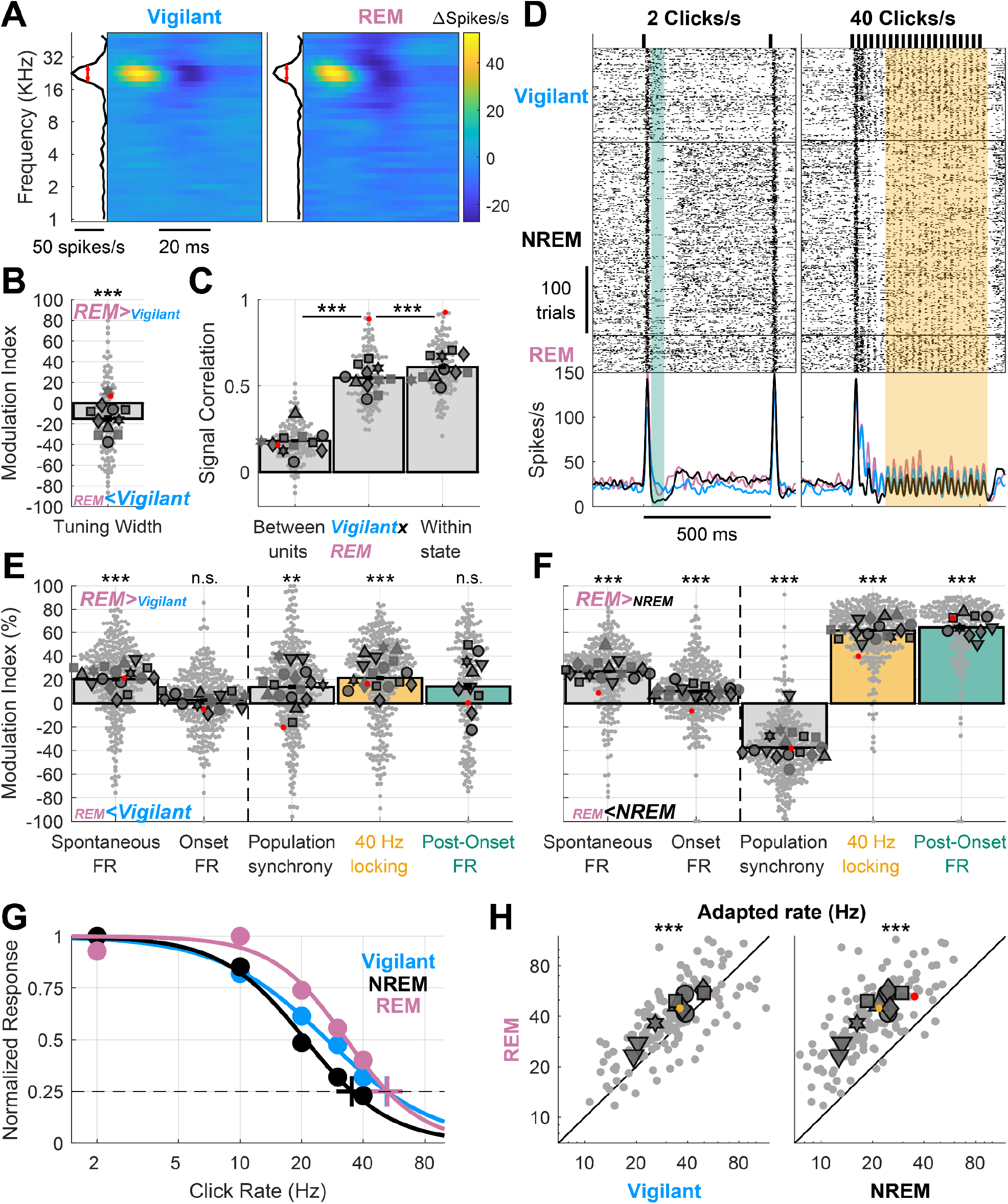
Auditory processing in REM sleep resembles wakefulness rather than NREM sleep. A) Representative spectro-temporal receptive field (STRF) of an auditory cortex unit showing preserved tuning during the Vigilant and REM sleep conditions (left and right, respectively). B) Modulation of frequency tuning width (REM sleep vs. Vigilant conditions) for all units (n=122) and sessions (n=11). C) Signal correlations of frequency tuning across the entire dataset between different units in the same session (left bar), between Vigilant and REM-sleep conditions of the same individual units (middle bar) and between 1st and 2nd halves of trials in the same condition for the same individual units (right bar). Note that signal correlations are nearly as high across Vigilant and REM-sleep conditions as they are within the same condition. D) An example unit raster and peristimulus time histogram (PSTH) for 2 and 40 clicks/s click trains (left and right, respectively). Green shading represents the post-onset [30,80]ms period and yellow shading represents the [130,530]ms period with sustained locking to the 40 click/s train. Note that in both these intervals, neuronal activity was similar in Vigilant and REM sleep, unlike the attenuation observed in NREM sleep. E) Modulation of spontaneous FR, onset response FR, population synchrony, 40-Hz locking and post onset FR during REM sleep relative to the Vigilant condition for all units (n=327/198) and sessions (n=17/10). 2 click/s train were only presented in 11 sessions (‘auditory paradigm A’). Most auditory processing features were comparable or enhanced in REM sleep compared with the Vigilant condition. Dashed vertical line separates features minimally/not significantly affected by NREM sleep/Tired as in previous figures, for reference. F) same as E but comparing REM sleep to NREM sleep. G) Normalized locked responses in a representative unit (y-axis) as a function of click rate (x-axis) separately for Vigilant (cyan), NREM sleep (green), and REM sleep (pink). Circles represent the observed locked response to each click rate in each condition. Thick traces represent the best sigmoid fit. Cross represents the estimated ‘adapted click-rate’, i.e. the click rate for which the normalized response would be 25% of maximum. H) Left: scatter plot of the ‘adapted click-rate’ for all units (n=138) and sessions (n=10), comparing REM sleep (y-axis) with Vigilant conditions (x-axis); Right: same when comparing REM sleep (y-axis) with recovery NREM sleep (x-axis). Yellow cross represents mean±SEM across all units. For Panels B, C, E, F and H: small gray markers represent individual units. Large dark gray markers represent mean of all units in an individual session. Each marker shape represents sessions from an individual animal. Markers with/without black edges represent ‘auditory paradigm A’ and ‘auditory paradigm B’ sessions, respectively. Red dots point to the representative unit presented in panels A and D.

## Discussion

The present results reveal how SD affects activity and stimulus-evoked responses in the auditory cortex. We find that some aspects of cortical auditory processing – including frequency tuning, spontaneous firing, and onset responses – are preserved across Vigilant and Tired conditions and are largely invariant to SD. By contrast, population synchrony, entrainment to fast click-trains, and post-onset silence are strongly modulated by SD (Fig. 2). The effects of SD on cortical auditory processing mimic those of NREM sleep, when similar effects manifest with stronger intensity (Fig. 3). Both SD and NREM sleep entail sensory adaptation at lower frequencies, suggesting that low-arousal states disrupt cortical processing of fast inputs (Fig. 4). We also find that stimulus-induced neuronal silent intervals are more sensitive to SD than are spontaneous silent intervals (‘OFF-states’, Fig. 5), a result that could been interpreted to show that perturbation reveals a hidden state of neuronal bi-stability not easily observed in spontaneous activity (Massimini et al., 2005; Vyazovskiy et al., 2009b). Finally, auditory processing during REM sleep (Fig. 6) resembles that in vigilant wakefulness (unlike NREM sleep) and highlights the key role of cortical desynchronization in auditory processing. Our results extend previous research showing that SD and drowsiness influences sensory processing (Kong et al., 2014; Muller-Gass and Campbell, 2019; Nir et al., 2017; Weissman et al., 2006; Wiggins et al., 2018) by showing that SD-induced changes already occur at primary cortices, earlier along the ascending cortical hierarchy than reported so far.

How do the present results stand with respect to whether primary cortices are robustly modulated, or largely invariant, to brain states and arousal? On one hand, the effects of states such as sleep and anesthesia are typically more modest in primary cortex than in high-order regions (Davis et al., 2007; Hayat et al., 2021; Krom et al., 2020; Liu et al., 2012; Makov et al., 2017; Nourski et al., 2018, 2016; Sela et al., 2020; Sellers et al., 2015; Sharon and Nir, 2018). Similarly, the effects of neuromodulation, attention, and consciousness are more prevalent in high-order regions compared to early sensory cortex (Atiani et al., 2014; Gelbard-Sagiv et al., 2018; Leopold and Logothetis, 1996). On the other hand, many studies report robust changes in early sensory cortex processing associated with arousal, task engagement, and other task parameters (Bagur et al., 2018; Banks et al., 2018; Carcea et al., 2017; Downer et al., 2015; Lin et al., 2019; Marguet and Harris, 2011; McGinley et al., 2015; Niwa et al., 2012; Otazu et al., 2009; Pachitariu et al., 2015; Sakata, 2016; Schwartz et al., 2020; Shimaoka et al., 2018; Zhou et al., 2014). Our results support a model in which specific features of the auditory response undergo increasing state-dependent deterioration along the sensory hierarchy. At the earliest processing stages - in peripheral sensory organs, thalamus, and primary cortices -response degradation gradually accumulates but on the whole is often modest and difficult to detect (Bereshpolova et al., 2011; Sakata, 2016; Scholvinck et al., 2015). Degradation builds up further along the cortical hierarchy, possibly due to higher sensitivity of inter-cortical signal transmission to behavioral states. Thus, in high-order regions, responses most correlated with perception often exhibit a sharper contrast between states. By focusing on responses in sensory cortex during SD, we were able to reveal state-dependent changes in specific features of neuronal response already at early auditory cortex.

Directly comparing different features (‘motifs’) of AC processing reveals which neural signatures are most sensitive to low-arousal. We find that SD and sleep only weakly affect neuronal tuning, spontaneous firing, and onset responses, compared with other aspects of auditory processing. The observation that frequency tuning is relatively invariant to SD and sleep is in line with the fact that it was traditionally studied successfully in anesthetized animals (Merzenich et al., 1975). However, while some studies report invariant tuning across states (Schwartz et al., 2020; Zhou et al., 2014), others report arousal-induced modulations in tuning (Gaese et al., 2001; Lin et al., 2019). Naturally, differences between separate studies can reflect changes in magnitude/type of arousal manipulation (e.g. sleep vs. anesthesia), species, cortical layer, or recorded cell types. The strength of the current study is that by comparing different motifs of auditory processing in the same neurons and experiments, our results provide important context in showing that frequency tuning is one of the most arousal-invariant feature of AC processing compared with other features we measured. We also find that SD and sleep only modestly affect baseline firing rates and onset response magnitudes in AC, in general agreement with previous reports showing modest changes during sleep (Issa and Wang, 2008; Nir et al., 2013a; Sela et al., 2020). While previous rodent studies reported increased spiking activity upon prolonged wakefulness and sleep deprivation (Fisher et al., 2016; Vyazovskiy et al., 2009a), we do not observe such increases, possibly due to our focus on early sensory cortex or due to differences in the sleep-deprivation method (Fisher et al., 2016).

By contrast to invariant features, population synchrony robustly increases upon SD (and even more so in NREM sleep), likely reducing the capacity of cortical circuits to represent information and support perception, consciousness, and behavior (Averbeck et al., 2006; Downer et al., 2015). Indeed, increased synchrony in neuronal populations at low frequencies (<20Hz) represents a core feature of low arousal states such as SD & sleep, spanning multiple levels from individual neurons, through circuits, to non-invasive global EEG recordings (Finelli et al., 2000; Nir et al., 2013b; Steriade et al., 1993; Vyazovskiy and Tobler, 2005; Vyazovskiy et al., 2011).

Our results extend previous work showing that reduced entrainment to fast inputs is a hallmark of unconscious low-arousal states. During deep anesthesia, responses to high-frequency stimuli are attenuated in cat visual cortex (Rager, 1998) and in rodent somatosensory (Castro-Alamancos, 2004) and auditory cortex (Marguet and Harris, 2011). In natural sleep and light propofol anesthesia, auditory cortex of both rodents and humans reveals reduced responses to 40Hz click-trains (Bergman et al., 2022; Hayat et al., 2021; Krom et al., 2020), as has been originally observed with scalp EEG (Lustenberger et al., 2017; Plourde, 1990). Here, we extend these results to show that already during wakefulness, SD-induced Tired conditions entail sensory adaptation at significantly lower frequencies, acting like a low-pass filter that quenches high-frequency neural inputs and diminishes rapid transmission of information across brain regions. The underlying mechanism may involve changes in short term synaptic plasticity, as the synaptic proteome was recently shown to be modulated by SD (Noya et al., 2019).

The most sensitive feature of auditory processing modulated by SD and NREM sleep is stimulus-induced neuronal silence, which has been suggested to reveal an underlying neuronal bi-stability in low-arousal states (Massimini et al., 2007). Such bi-stability may not allow neurons in low-arousal states to maintain sustained firing in response to a stimulus, and its occurrence in some cortical regions may underlie the behavioral inability to successfully maintain sustained attention (Vyazovskiy et al., 2011). While we cannot definitively demonstrate that stimulus-induced silent intervals reflect genuine membrane potential bi-stability (Up and Down states), we believe our results agree with that interpretation. For one, the fact that silent intervals don’t appear after vigorous spontaneous spiking (Fig. 5A-C), strengthens the notion that stimulus-induced silent intervals indeed reflect a network level phenomenon, not just individual neurons showing suppressed FR after vigorous spiking. Importantly, stimulus-induced activity reveals a hidden facet of neuronal activity during SD (propensity for silent intervals) that is not readily observed in spontaneous activity (Vyazovskiy et al., 2013). Our results join previous work with transcranial magnetic stimulation (TMS) in humans (Massimini et al., 2007), as well as electrical intracortical stimulation in rodents (Vyazovskiy et al., 2013, 2009b), both showing that perturbation can reveal the latent state of cortical activity and trigger a slow wave at any time during NREM sleep, even when the ongoing EEG shows little spontaneous slow wave activity. Indeed, quantifying the brain’s response to perturbation (e.g. with TMS-EEG) offers a more sensitive approach to detect bi-stability that accompanies disorders of consciousness and brain-injured patients (Casali et al., 2013). Our results suggest that examining the response to sensory stimuli might be a particularly effective way to assess the level of drowsiness and sleep deprivation (for example in the context of human EEG during driving).

Our results extend the notion that NREM-sleep-related activities invade the activity of the waking brain after SD. This has been established for spontaneous EEG activity (‘EEG slowing’) (Finelli et al., 2000; Nir et al., 2017; Vyazovskiy and Tobler, 2005) and for ongoing neuronal activity and OFF states (Vyazovskiy et al., 2011). Here we show that SD mimics NREM sleep also in how it affects sensory processing, and specifically in early sensory cortex.

In contrast to NREM sleep, REM sleep resembles vigilant wakefulness for all features of cortical auditory activity and stimulus-evoked responses. REM sleep serves as a unique test-case to determine which elements of neuronal activity and sensory responses reflect disconnection from environmental sensory stimuli vs. elements that reflect the ability of the brain to generate conscious experience (whether externally- or internally-generated). On one hand, REM sleep is similar to NREM sleep in that both entail disconnection from the external world; on the other hand, REM sleep is similar to vigilant wakefulness in that during both states the brain generates conscious experience. Thus, the result that AC activity in REM sleep resembles vigilant wakefulness suggests that the changes in cortical auditory processing observed in SD and NREM sleep may reflect features of an unconscious brain state, and that sensory disconnection can co-occur with desynchronized wake-like processing in AC. These results point to a key role for cholinergic modulation in AC processing, given that high acetylcholine levels drive cortical desynchronization similarly across REM sleep and wakefulness (Nir and Tononi, 2010). Future studies could directly study whether cholinergic modulation of auditory pathways is necessary and sufficient to support specific features of auditory processing as observed in vigilant wakefulness.

Some limitations of the study should be explicitly acknowledged. First, our procedure for implanting microwire arrays did not enable us to obtain reliable information about the cortical layer and type of recorded neurons. Thus, our sample could be biased and best capture specific subpopulations such as large pyramidal cells with higher baseline firing rates that register more readily in extracellular recordings. Second, the state-dependent changes observed in early auditory cortex could possibly be inherited from earlier regions such as the auditory thalamus, not recorded here. Third, our data from NREM and REM sleep reflects recovery sleep, likely associated with deeper sleep and stronger attenuation than usual.

Still, the fact that robust changes in auditory processing were observed during SD while the animal was awake, and no such changes were observed during REM sleep, partly alleviates that concern. Fourth, as no behavioral task was included in the study it remains to be seen if the reported changes in auditory processing are associated with the deterioration in behavioral performance (‘lapses’) typical of SD. Future studies could examine if moment-to-moment variability in behavioral performance is associated with moment-to-moment changes in sensory processing. Finally, another important aspect to address is the possibility that SD periods were significantly contaminated by brief sleep episodes, which in turn may have driven the changes in auditory processing seen in the Tired condition. We don’t believe this is the case, since video monitoring did not reveal periods of sleep during SD. In addition, EEG slow wave power during the Tired condition was largely comparable to the Vigilant condition, but very different from that during NREM sleep.

In conclusion, we examined the effects of SD and recovery sleep on different aspects of auditory cortex processing and found that SD already affects neural processing in early sensory cortex. We find that SD robustly modulated some aspects of auditory processing (population synchrony, entrainment to fast inputs, and stimulus-induced silent intervals) while other aspects remained stable (neuronal tuning, spontaneous firing and onset responses). Stimulus-induced activity reveals a hidden aspect of neuronal bi-stability that is not observed in spontaneous activity. This is important both conceptually and for practical/clinical applications, as it offers new ways to monitor sleepiness with greater sensitivity. Finally, changes in auditory processing during SD are qualitatively similar to those observed during NREM sleep but not REM sleep, suggesting that NREM-sleep-like processes are specifically invading activity of the waking brain in SD and disrupt behavior.

## Methods

### Animals

Experiments were performed in seven male Wistar rats individually housed in transparent Perspex cages with food and water available ad libitum. Ambient temperature was kept between 20°-24° Celsius and a 12:12 hours light/dark cycle was maintained with light onset at 10:00 AM. All experimental procedures, including animal handling, sleep deprivation and surgery, followed the National Institutes of Health’s Guide for the care and use of laboratory animals and were approved by the Institutional Animal Care and Use Committee of Tel Aviv University.

### Surgery and electrode implantation

Prior to surgery, microwire arrays were coated with a thin layer of DiI fluorescent dye (DiIC18, Invitrogen) under microscopic control to facilitate subsequent localization. Surgery was performed as previously described (Sela et al., 2020). First, induction of general anesthesia was achieved using isoflurane (4%). Animals were then placed in a stereotactic frame (David Kopf Instruments) and maintained for the rest of the surgery under anesthesia (isoflurane, 1.5-2%) and 37°C body temperature (closed-loop heating pad system, Harvard Apparatus). Animals were administered antibiotics (Cefazolin, 20 mg/kg i.m.), analgesia (Carpofen, 5 mg/kg i.p.) and dexamethasone (0.5 mg/kg, i.p.). Their scalp was shaved and liquid gel (Viscotears) was applied to protect the eyes. lignocaine (7 mg/kg) was infused subcutaneously before incision and then the skull was exposed and cleaned. Two frontal screws (one on each hemisphere, 1mm in diameter) and a single parietal screw (left hemisphere) were placed in the skull for recording EEG. Two screws, serving as reference and ground, were placed above the cerebellum. Two single-stranded stainless-steel wires were inserted to the neck muscles to record EMG. EEG and EMG wires were soldered onto a head-stage connector (Omnetics). Dental cement was used to cover all screws and wires. A small craniotomy was performed over the right hemisphere, and the dura was carefully dissected. A 16-electrode microwire array targeting the auditory cortex was implanted (Tucker-Davis Technologies, TDT, 33 or 50µm wire diameter, 6-6.5 mm long, 15° tip angle; arrays consisting of 2 rows × 8 wires, with 375µm medial-lateral separation between rows and 250µm anterior–posterior separation within each row). Implantation was diagonal (angle of 28°, see Fig. 1B) using insertion point center coordinates of P: - 4.30mm, L: 4.5mm relative to Bregma, and inserted to a final depth of 4.6mm. Following implantation, a silicone gel was applied to cover the craniotomy (Kwik-Sil; World Precision Instruments) and Fusio (Pentron) was used to fix the microwire array in place. At the end of the surgery, chloramphenicol 3% ointment was applied topically and additional analgesia was provided by injecting buprenorphine systemically (0.025 mg/kg s.c.) as the rat awoke from anesthesia. Dexamethasone (1.3 mg/kg) was given with food in the days following the surgery to reduce pain and inflammation around implantation.

### Histology

Upon completion of the experiments, position of electrodes was verified by histology in 4 out of 7 animals (e.g. Fig. 1B). Animals were transcardially perfused with 4% paraformaldehyde (PFA) under deep (5% isoflurane) anesthesia. Brains were refrigerated in PFA for a week, cut into 50–60µm serial coronal sections using a vibrating microtome (Leica Biosystems), and stained with fluorescent cresyl violet/Nissl (Rhenium). Histological verification confirmed that electrodes were located within areas Au1/AuD as defined by (Paxinos and Watson, 2006).

### Electrophysiology

As previously described in (Sela et al., 2020), data was acquired using a RZ2 processor (TDT) with microwire extracellular activity digitally sampled at 24.4 kHz (PZ2 amplifier, TDT) and EEG and EMG pre-amplified (RA16LI, TDT) and digitally sampled at 256.9 Hz (PZ2 amplifier, TDT). Spike sorting was performed using “wave_clus” (Quiroga et al., 2004), employing a detection threshold of 5 SD and automatic superparamagnetic clustering of wavelet coefficients. Clusters were manually selected, refined, and tagged as multi- or single-unit based on stability throughout recording, quality of separation from other clusters, consistency of spike waveforms and inter-spike interval distributions as in (Nir et al., 2013a).

### Experimental Design

In the week preceding the surgery, subjects were habituated to spending time inside the motorized running wheel for a few hours every day (Fig 1A), and then gradually to participating in the sleep deprivation protocol (Fig. 1D, see below).

We ran 19 sleep deprivation experimental sessions, as follows. At light onset (10 AM) rats were moved from their home cage to a motorized running wheel (Fig 1A, Model 80860B, Lafayette Instrument) placed inside a sound-attenuation chamber (−55dB, H.N.A) and underwent 5 hours of sleep deprivation. Throughout the sleep deprivation period, the wheel was intermittingly slightly rotated for 3 seconds, forcing a short running bout, with a randomly chosen 12-18 seconds interval break in between running bouts. Next, rats were left undisturbed in the fixed wheel for a recovery sleep opportunity period of 5 hours. Auditory stimulation (below) was delivered intermittently throughout each session, during both sleep deprivation and recovery sleep periods, without regard to the wheel’s movement regime.

### Auditory stimulation

Sounds were synthesized in Matlab (MathWorks) and transduced into voltage signals by a high-sampling rate sound card (192 kHz, LynxTWO, Lynx), amplified (SA1, TDT) and played free-field through a magnetic speaker (MF1, TDT), mounted 60 cm above the motorized running wheel. We employed two different auditory paradigms on separate sessions/days:

#### Auditory paradigm A

(11 Sessions, 7 animals, markers with black edges accompanying histograms in figures e.g. Fig. 2B): Stimuli included click trains and a set of Dynamic Random Chords (DRCs, (Linden, 2003)). Click trains were 500ms in duration at rates of either {2, 10, 20, 30, 40} clicks/sec. DRCs were 2.5s in duration and included a train of randomly chosen 20ms “chords”, each comprised of an average of 6 randomly chosen tone-pips at different frequencies (1-64 KHz, with 1/6 octave intervals, 5ms cosine ramp, fixed sound level). There were 190 different DRC stimuli. A typical 10h session contained 2000 blocks, each consisting of a single DRC stimulus and a single repetition of each click train (presented at random order), and with an inter-stimulus interval of 2s and ±0.25s jitter.

#### Auditory paradigm B

(8 Sessions, 6 animals, markers without black edges accompanying histograms in figures e.g. Fig. 2B): Stimuli included a 40 clicks/s click-train, and a different set of DRC stimuli with denser sampling of the frequency and intensity axes (better resolution) to allow for quantitative assessment of neuronal tuning curves. We used 6s trains of randomly chosen 20ms “chords”, each comprised of an average of 12 randomly chosen tone pips at different frequencies and different sound levels (1-64 KHz, with 1/10 octave intervals, 5ms cosine ramp, spanning an 80 dB range in 10 dB intervals). There were 120 different DRC stimuli. A typical 10h session contained 600 blocks, each consisting of a single DRC and 4 repetitions of the 40 Hz click train, presented at random order, with an inter-stimulus interval of 2s and ±0.25s jitter.

Both paradigms included an 8s inter-stimulus interval every 2 minutes.

### Sleep scoring and analysis of arousal states

Manual sleep scoring was performed offline for the entire experimental session, employing visual inspection of EEGs, EMGs and video/behavior as in previous studies (Nir et al., 2013a; Rodriguez et al., 2016; Sela et al., 2020; Vyazovskiy et al., 2011). First, we excluded any periods when the wheel was moving (forced running bouts during sleep deprivation) and other periods of active wakefulness with behavioral activity (e.g., locomotion, grooming) as confirmed with video. Next, we categorized periods to either wakefulness (low-voltage high-frequency EEG activity and high tonic EMG with occasional phasic activity), NREM sleep (high-amplitude slow wave activity and low tonic EMG activity), REM sleep (low-amplitude wake-like frontal EEG co-occurring with theta activity in parietal EEG and flat EMG), or unknown periods not analyzed further (e.g. state transitions, to conservatively remove these epochs for subsequent analysis).

Next, each auditory stimulation trial was categorized to one of four conditions: Vigilant, Tired, NREM and REM, as follows. Vigilant and Tired categories comprised of the first or last third of (quiet) wakefulness trials during the sleep deprivation period, respectively, while NREM and REM comprised of trials scored as such during the recovery sleep period. To assert that differences between the Vigilant and NREM sleep categories did not stem from temporal order effects (e.g. Vigilant trials always preceding NREM by a few hours), we also defined a fifth condition – quiet wakefulness during the recovery sleep period, denoted as QW-RSP. Neural activity during QW-RSP was very similar to the Vigilant condition earlier in the experiment, qualitatively replicating the results of differences between Vigilant and NREM conditions (data not shown).

### Analysis of auditory responses across states

#### Neuronal Tuning analysis

(Fig. 2A-C, 3A-C, 6A-C). To analyze responses to the two sets of DRC stimuli (Paradigms A and B) we performed the following analysis. Given that tone pips at each frequency were presented independently (statistically), we calculated the effects of each tone-pip on neuronal firing rates as: 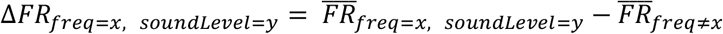. Tuning width (Fig. 2B, 3B, 6B) was calculated as the Full-Width Half Maximum (FWHM) around the best frequency in octaves (red lines in Fig. 2A, 3A, 6A). In paradigm B, frequency tuning width was calculated for the loudest sound level. The tuning width Modulation Index (MI) between any two conditions was defined as (and similar to Gain Index in (Sela et al., 2020)): 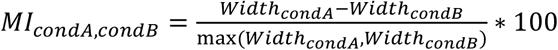

Due to a technical problem in the presentation of tones at the highest frequency of 59.7kHz, many units exhibited maximal responses to this particular frequency, so these trials were removed from subsequent analysis to ensure result validity. To calculate the signal correlation of the neuronal tuning between any two conditions we conducted the following analysis (Fig. 2C, 3C, 6C): in paradigm A, where there was only a single sound level, the neuronal tuning map was defined as the spectro-temporal receptive field (STRF, see spectrograms in Fig 2A, 3A, 6A) a F×T matrix (where F is number of frequencies-[1,64] KHz with 1/6 octave steps, and T is the number of time points [0,50]ms) with each value representing the ΔFR (above baseline) for a given frequency and time-point. The STRF map was smoothed in the temporal domain with a Gaussian kernel (σ=5ms). In paradigm B the tuning map was defined as the frequency response area (FRA) a F×L matrix (where F is number of frequencies-[1,64] KHz with 0.1 octave steps, and L is the number of sound levels [0,80] dB in 10dB steps), with each value representing the ΔFR (above baseline) for a given frequency and sound-level (in the [5,30]ms temporal window). The FRA map was smoothed in the frequency domain with a square window (length=0.3 octaves). The signal correlation between any two conditions is defined a point-by-point Pearson correlation between the two conditions tuning maps (STRF for paradigm A, and FRA for paradigm B). Realistically however, this correlation will always be smaller than one, since the neural response inevitably contains some noise, and because estimates of the response are limited by a finite number of trials. The signal correlation is also expected to be on average larger than zero, as even different units in the same region might show similar preference to frequency and temporal profile, yielding positive signal correlation. Therefore, to create meaningful benchmarks to compare signal correlations, we compared the following three values for each unit separately: (i) [minimal correlation expected]: signal correlation of each neuron’s tuning map (STRF/FRA for paradigms A/B, respectively) with the tuning maps of *other units* in the session across different conditions (left bar in Fig. 2C, 3C, 6C), (ii) [main value of interest]: signal correlation of each neuron’s tuning map in one condition (e.g. Vigilant) with its tuning map in the other condition (e.g. Tired, middle bar in Fig. 2C, 3C, 6C), (iii) [maximal possible correlation]: each neuron’s signal correlation of its tuning map in the 1^st^ vs. 2^nd^ half of trials in the same condition (right bar in Fig. 2C, 3C, 6C). Formally:

{u_1_, u_2_, …, u_n_} a set of n Units in a given session.

{s_1_, s_2_} a set (S) of two Conditions we want to compare (e.g. Vigilant and Tired).

{h_1_, h_2_} first and second half of trials for a given condition.

*TM*_*u,s,h*_ is the Tuning-Map (STRF/FRA matrix for paradigms A/B, respectively) of Unit u for h half of trials in Condition s.

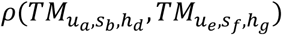 is the point-by-point Pearson correlation coefficient between the two tuning map matrices 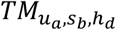 and 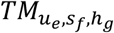.

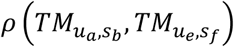, the correlation between the tuning maps of unit u_a_ in condition s_b_ and unit u_e_ in condition s_f_ is defined as mean correlation coefficient between all halves combinations:

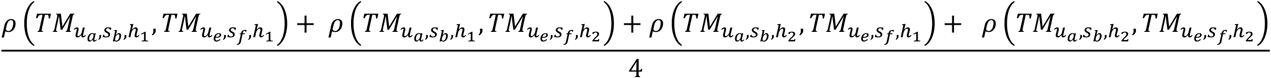

The three different measures of signal correlation (left, middle and right bars, respectively) for a given neuron u_i_ are defined as:

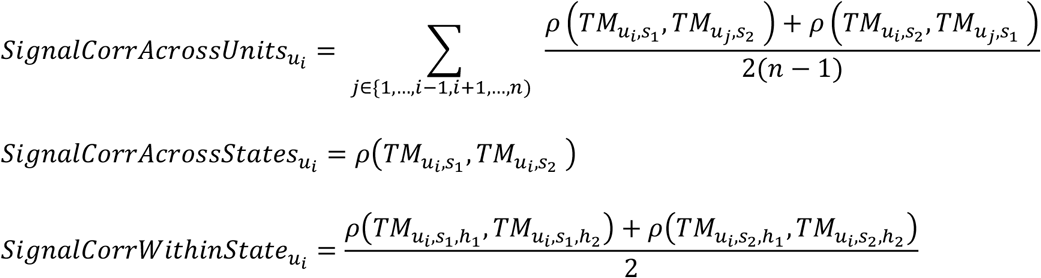

#### Analysis of responses to click trains

(Fig. 2D-E, 3D-E, 6D-F). Spontaneous firing rate (FR) was calculated as the mean firing rate in the [-500,0]ms window preceding the click-trains stimuli, and post-onset FR as the mean FR in the [30,80]ms window. Onset response and sustained locking to different click rates (Fig. 2D-E, 3D-E, 4A-C, 6D-H) were obtained from the smoothed peri-stimulus time histogram (PSTH, Gaussian kernel, s=2ms). Onset response was obtained by extracting the maximal firing rate during the [0,50]ms window of the smoothed PSTH. Locking to different click rates (2, 10, 20, 30 & 40 clicks/s) was obtained by calculating the mean firing rate for each phase during the inter-click intervals in the [130,530]ms window. Then, firing rate locking was defined by the minimum firing rate (during the least preferred phase relative to the click) subtracted from the maximum firing rate (during the most preferred phase). Population synchrony was defined as population coupling (Okun et al., 2015), the correlation of each unit firing to that of the entire neuronal population average in 50ms bins during baseline ([-1000,0]ms). Population coupling was calculated for each trial baseline period and then averaged for all trials in a given condition.

Modulation index between two conditions for all measures above was calculated as for the tuning width 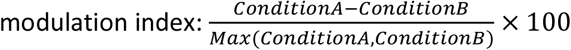

#### Sensory adaptation curve fitting

(Fig. 4D-F, 6I-J). We first normalized each unit’s sustained locking response to each click rate by dividing its firing rate to the maximum of all locked responses across all rates (2,10,20,30,40 clicks/s) and its onset response during the same condition (points in figure 4D and 6I). we then fitted the data (the five normalized responses: 2,10,20,30,40 clicks/s) with the following sigmoid model, where x_0_ is the click-rate where the normalized response is 0.5 (50% of max) and k is the slope of decay of the response.

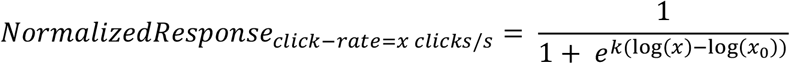

Using this fitted model (traces in Fig. 4D, 6I) we estimate for each neuron in each condition the ‘adapted rate’, defined as the estimated click rate for which the normalized response would be 0.25 (25% of the maximum, crosses in Fig. 4D, 6I; calculation with other percentile cutoff of maximum did not affect the results). In examining how the adapted rate changes across different conditions for the entire neuronal population (Fig 4E, 6I) and in an effort to exclude noisy responses, we included in the population analysis only units with satisfactory sigmoid fit (rms<0.07) and for which the adapted click rate was within the range of [2,150] clicks/s. This criterion led to the exclusion of a minority of neurons (47/197 units, 23.9%).

#### Silent intervals analysis

(Fig. 5). To consider the effects of sleep deprivation and NREM sleep on spontaneous and stimulus-induced silent intervals we performed the following analysis. We created a raster plot of spontaneous spiking bursts for each unit (Fig. 5A left), which was trial-by-trial matched to the its click-induced onset response (Fig. 5A right). This was done by matching each trial of 2-Hz click-train onset response ([0,30]ms) with an identical (or as similar as possible) spike train obtained during spontaneous activity in the same arousal condition. Each unit PSTH was normalized to its baseline FR and a grand-mean PSTH was calculated for stimulus-induced responses and matched spontaneous bursts (Fig. 5B). We quantified the effect per unit by calculating the mean baseline-normalized FR in the post-onset temporal window ([30,80]ms) for each condition (Vigilant, Tired and NREM sleep) and for stimulus-induced and spontaneous spiking bursts.

To detect (possibly local) silent intervals we performed our analysis on a per-microwire basis (aggregating the spikes from all clusters recorded in the microwire). We defined silent intervals as periods of 50ms neuron silence (Vyazovskiy et al., 2009a) and checked their probability in the baseline ([-50,0]ms), as well as post-onset ([30,80]ms) period (spontaneous and stimulus-induced silent intervals in Fig. 5, respectively, orange and green lines). To control for changes in silent interval probability stemming simply from changes in the spontaneous firing rate, we took the absolute silent interval probability and subtracted the expected silent interval probability from a simulated Poisson-process unit activity with the same firing rate (Δ50ms silence probability in Fig. 5E). To formally compare the effects of sleep deprivation on spontaneous vs. stimulus-induced silent interval probabilities (Fig 5C) we calculated the following modulation index:

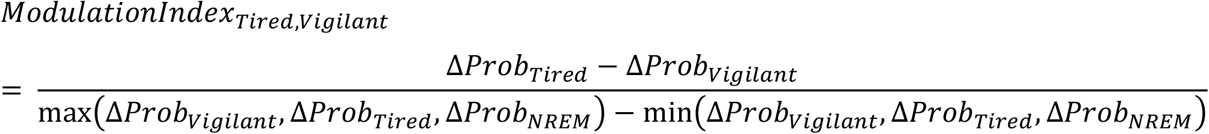

### Statistics

Due to the nested and hierarchical nature of electrophysiological neural data (Aarts et al., 2014; Makin and Orban de Xivry, 2019) we used a linear mixed-effects model (LME). The LME was used to account for non-independencies in measures from different units that were obtained in the same electrode, experimental session or animal. Animal identity was used as a random effect, together with experimental session and microwire electrode as nested random effects within each animal. Model parameters were calculated using ‘fitlme’ function (Matlab, MathWorks) using restricted maximum likelihood estimation. In the cases where the data samples were obtained on a per-microwire basis (analysis in Fig. 5D-F, instead of per-unit basis) only animal identity and experimental session (nested within animal) were used as random effects. Using conservative non-parametric statistical tests (Wilcoxon Rank-Sum Test or Wilcoxon Sign-Rank Test) on the data summarized at the level of animals (n=7) or sessions (n=19) yielded qualitatively very similar results in terms of statistical significance as the LME model (data not shown). In figures depicting mean effects per session (large markers in figures 2B,2C, 2E, 3B, 3C, 3E, 4B, 4C, 4E, 5C, 5E, 5F, 6B, 6C, 6E, 6F and 6H) only sessions with at least 5 units were included. The LME analysis however, was always applied on all sessions, even those with few units. If not stated otherwise all effect sizes mentioned in main text are described as mean±SEM over all units. When testing for variance across multiple (>2) conditions a Friedman test was used (akin to a non-parametric repeated measures ANOVA) on data summarized at the level of animals (n=7, averaging all the units for each animal).

## Acknowledgements

We thank Israel Nelken for input on auditory setup and analyses. Rafael Malach, Mark Shein-Idelson, Aaron Krom, Yael Oran and Yaniv Sela for comments on an earlier draft, and members of the Nir Lab for discussions.

---

**Supplementary Figure 1 (relates to Fig. 2).**
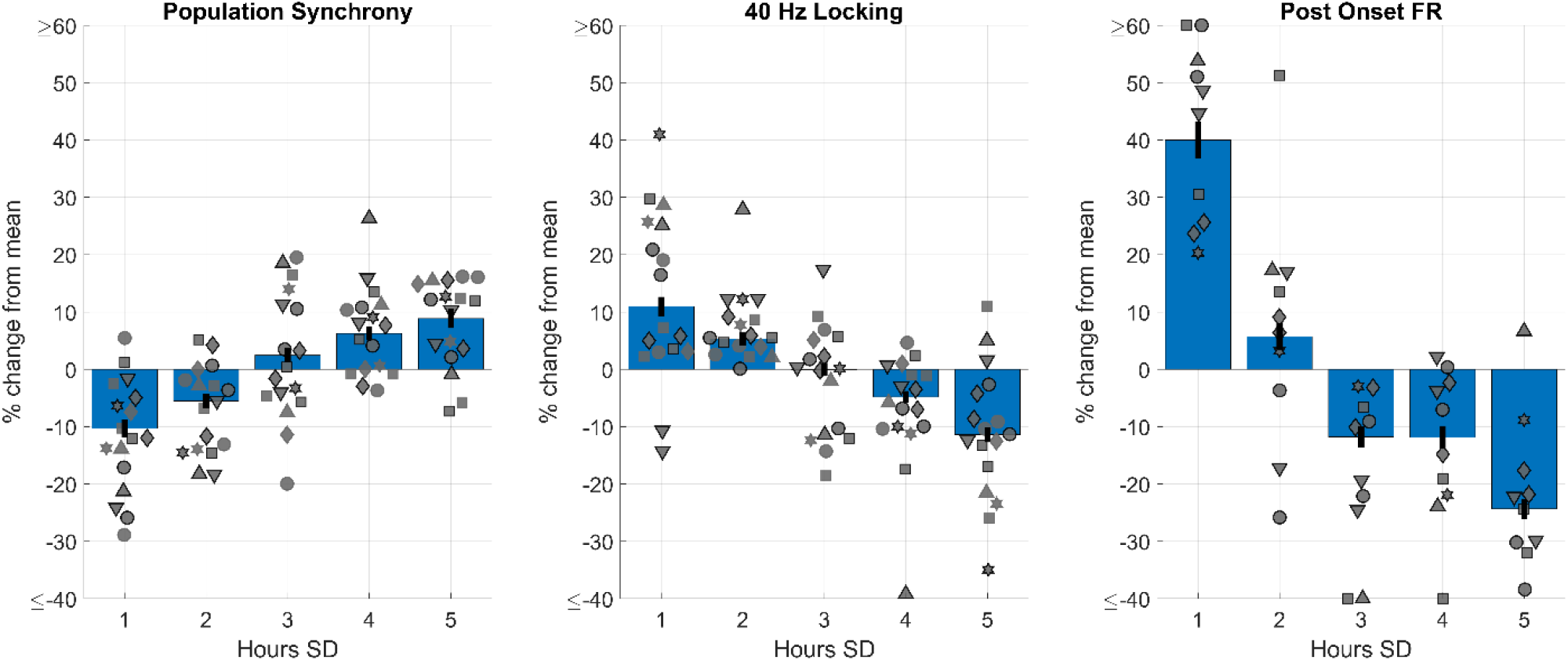
Gradual changes in SD-sensitive ‘motifs’. Mean % changes at 1-hour time bins (20% of trials) during the SD period for three different sensitive neural processing ‘motifs’: population synchrony (left), 40-Hz click train locking (middle) and post-onset FR (right). Individual markers depict mean % change of all units is a single session. Different marker shapes represent different animals. Bars and black lines depict the mean and SEM across all sessions, respectively.

**Supplementary Figure 2 (relates to Fig. 4).**
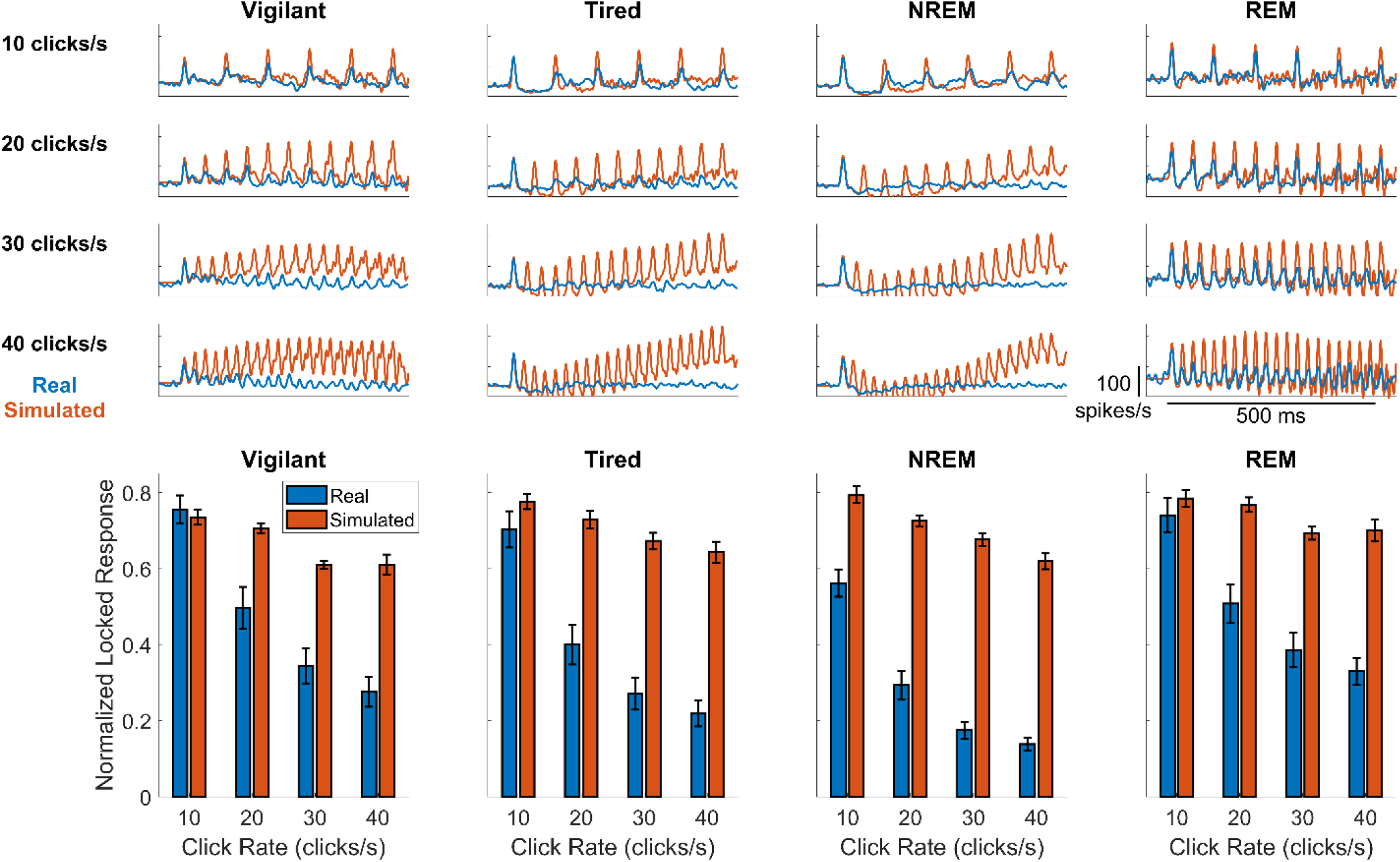
Post-onset FR reduction doesn’t explain reduced locking to rapid click trains. We examined if decreased locking to rapid click trains may be trivially explained by post-onset FR suppression that may coincide with the evoked response to subsequent clicks. We constructed a simple linear model aiming to predict the response to different click trains by shifting in time and summing up the average response to an individual click (Methods). Top) an example of individual unit locked response to different click rates (rows) across different conditions (columns). Blue traces represent the actual response while red traces represent the linear model. For this unit the model predicts much stronger locking to fast click trains than that is observed in practice (compare red to blue traces at the bottom row). Bottom) mean normalized locked response for different conditions (columns) and click rates (different bars). Blue and red bars represent the mean real and modeled response across all units, respectively. The large gap for fast click trains (especially for NREM and Tired conditions) demonstrates that post-onset FR reduction seen in response to individual clicks doesn’t trivially explain reduced locking to fast click trains.

